# CHEK2 SIGNALING IS THE KEY REGULATOR OF OOCYTE SURVIVAL AFTER CHEMOTHERAPY

**DOI:** 10.1101/2021.09.23.461589

**Authors:** Chihiro Emori, Zachary Boucher, Ewelina Bolcun-Filas

## Abstract

Radiation and chemotherapy can damage the primordial follicle reserve in female cancer patients leading to ovarian failure and infertility. Preservation of ovarian function requires treatment strategies that prevent loss of immature oocytes in primordial follicles during cancer therapy. Checkpoint kinase 2 (CHEK2) inhibition prevents loss of primordial oocytes caused by DNA damage and thus is a promising target for ovoprotective treatment against genotoxic agents. To determine which cancer treatments could benefit from ovoprotective activity of CHEK2 inhibition we investigated oocyte survival in *Chek2*-/- mice exposed to different chemotherapy drugs. Here, we show that loss of CHEK2 function prevents elimination of primordial oocytes damaged by cisplatin, cyclophosphamide, mafosfamide, doxorubicin, and etoposide, suggesting it could be used to reduce ovarian damage caused by wide range of drugs. Using genetic knockouts we reveal a critical role for TRP53 in oocyte response to chemotherapy drugs and show that both targets of CHEK2—TAp63 and TRP53— are activated by cisplatin and cyclophosphamide. Furthermore, we show that checkpoint kinase inhibitor and radiation- and chemotherapy sensitizer AZD7762 reduces oocyte elimination after radiation and chemotherapy treatments, despite its cytotoxic effect on ovarian somatic cells. Altogether, these findings demonstrate the role for CHEK2 as the master regulator of primordial oocyte survival or death and credential its targeting for ovoprotective treatments.

**SIGNIFICANCE:** Chemotherapy and radiation are ovotoxic and increase the risk of premature ovarian insufficiency and infertility in women cancer survivors. Development of treatment strategies preserving ovarian function and ensuring future reproductive health of female cancer patients depends on better understanding of the mechanisms underlying ovarian toxicity caused by different chemotherapy treatments. Preservation of long-term ovarian function can only be achieved by preventing the loss of immature oocytes in primordial follicles during toxic cancer therapies. Checkpoint kinase 2 (CHEK2) inhibition is an attractive strategy for protecting ovarian reserve with a potential additional benefit of sensitizing cancer cells to radiation and chemotherapy. Using a genetic approach, we show that blocking CHEK2 function is sufficient to prevent elimination of primordial oocytes damaged by chemotherapy drugs such as cisplatin, cyclophosphamide, mafosfamide, doxorubicin and etoposide. Many chemotherapy drugs are used in combination (e.g. cyclophosphamide with doxorubicin), thus the protective effect of CHEK2 inhibition is likely to be beneficial for a broad spectrum of patient treatments.

## INTRODUCTION

Women are born with a finite supply of the oocytes that ultimately form the eggs that are ovulated. The primordial oocytes are enclosed in primordial follicles (PMF) which must survive and last through a woman’s entire reproductive life. PMFs decrease naturally with age until menopause, which occurs when follicle supplies become depleted. Studies in human and animal models show that various environmental factors—such as pollutants, smoking, diet, etc.—can diminish ovarian follicle reserve (1–4). Importantly, now that diagnosis and treatment of cancers is more efficient, the ovarian follicle reserve can also be damaged in females patients by cancer treatments leading to accelerated PMF loss and resulting in infertility and premature ovarian insufficiency (POI) (1,5–7). Treatment-induced-POI results in early menopause, an endocrine deficiency associated with osteoporosis, cardiovascular disorders, depression, and other co-morbidities that require medical attention and can negatively impact the quality of survivorship (8–11). Quality of life considerations are especially important for pediatric, adolescent, and young adult cancer survivors due to their long-life expectancy. The overall prevalence of POI varies considerably (6-80%) due to heterogeneity of cancers and treatments (9,12–17). Fertility preservation methods, such as cryopreservation of eggs and embryos, are available for cancer patients of reproductive age; however, these methods are not available to prepubertal girls (18–20). Currently, ovarian tissue cryopreservation is the only fertility preservation method for prepubertal girls, but it does not prevent POI (21–24). There are currently no proven methods to protect ovaries from systemic chemotherapies during ongoing treatments in young patients.

Genotoxic cancer treatments kill cancer cells by inducing DNA damage, which is more detrimental to fast dividing cancer cells than healthy cells. However, these treatments can also damage healthy cells including oocytes. DNA damage inflicted in oocytes in PMFs seems to be the major trigger of radiation- and chemotherapy-induced PMF elimination (25–27); therefore, a strategy blocking the DNA damage response (DDR) in oocytes offers an attractive strategy to prevent PMF loss and POI. In contrast, growing oocytes in primary and secondary follicles are not as susceptible to DNA damage (28). This suggests a different response mechanism in the growing oocyte after follicle activation or a role for supporting granulosa cells within the follicle.

Why primordial oocytes, which are arrested at meiotic prophase I and are not replicating, are sensitive to chemotherapy drugs that interfere with DNA replication and transcription is still poorly understood. The most lethal type of DNA damage—DNA Double Strand Breaks (DSB)—leads to activation of CHEK2. CHEK2 coordinates DDR through activation of TRP53 (also known as p53) which leads to cell cycle arrest, senescence, or apoptosis depending on the cell type and cell cycle phase (29–31). In contrast, in primordial oocytes, CHEK2 predominantly activates a TRP53-related protein TRP63—and specifically its TA isoform (henceforth TAp63)—leading to oocyte elimination (26,28,32). Genetic inactivation of TAp63 protects primordial oocytes from cisplatin-induced damage but not from cyclophosphamide (33,34). These differences in survival suggest involvement of other DDR proteins (i.e., CHEK1) in the primordial oocyte response to different chemotherapy drugs. Studies utilizing kinase inhibitors known to target both CHEK1 and CHEK2 reported their protective effects in primordial oocytes against cisplatin, cyclophosphamide, and doxorubicin toxicity (32,34,35). However due to concerns about target specificity of these inhibitors it remains unclear which of the CHEK kinases responds to each chemotherapy drug. Considering the potential use of CHEK2 inhibitors for primordial oocyte protection, more information is needed regarding CHEK2 and its downstream effectors in chemotherapy-induced oocyte loss to identify cancer treatments which could benefit from oocyte-protective activity of CHEK2 inhibitors. Furthermore, better understanding of the cellular damage and related response mechanisms leading to primordial oocyte death will provide insight into the maintenance of the ovarian reserve in women exposed to other exogenous ovotoxic agents.

Our previous studies showed that genetic inactivation of CHEK2 prevents PMF loss in mice after exposure to ionizing radiation (26). In this study, we use wildtype and CHEK2-deficient mice to determine the role of CHEK2 in oocyte elimination in response to chemotherapy-induced damage and if inactivation of CHEK2 or its downstream effectors is sufficient to prevent chemotherapy-induced loss of PMFs. We provide genetic evidence that suggests CHEK2 inhibition is sufficient to protect primordial oocytes from alkylating agents cisplatin (CDDP) and cyclophosphamide (CTX), and topoisomerase II poisons doxorubicin (DOX) and etoposide (ETO). Moreover, we present genetic evidence that both TAp63 and TRP53 are involved in elimination of primordial oocytes damaged by CTX and CDDP. We also show that cancer sensitizing agent and CHEK1/2 inhibitor AZD7762 can protect oocytes from radiation and chemotherapy induced damage. The results of this study highlight the importance of targeted development of new inhibitors specific to CHEK2 to prevent PMF loss during cancer treatments.

## RESULTS

### Inactivation of CHEK2 prevents oocyte elimination caused by chemotherapy drugs in *ex vivo* organ culture

Previous studies showed that genetic and pharmacological inactivation of CHEK2 prevents POI in mice after exposure to radiation (26,36). Additional studies using the CHEK2 inhibitor BML-277 showed a protective effect against the chemotherapy drugs CDDP, DOX, and CTX derivative 4-HC (32,34,35). However, it remains unclear whether this protective effect is due to inhibition of CHEK2 or CHEK1 kinases due to concerns about inhibitor specificity (34). To test the direct role of CHEK2 in elimination of PMFs damaged by genotoxic chemotherapy drugs and determine if CHEK2 inhibition would be effective and sufficient in reducing PMF loss, we tested how genetic ablation of CHEK2 function affects primordial oocyte survival after treatments. We chose chemotherapy drugs known to induce DNA damage via different mechanisms and characterized by different severities of ovarian adverse effects: DNA alkylating agents CDDP and mafosfamide (MAFO; an analog of CTX (37)), and DNA topoisomerase II poisons DOX and ETO (33,38–41). We used MAFO in *ex vivo* culture to imitate the *in vivo* activity of CTX, which needs to be metabolized by the liver to generate metabolites with alkylating properties (37,42). To better control for effective drug doses and duration of exposure for comparative analysis we utilized *ex vivo* organ culture system, where we cultured whole ovaries to prevent potential PMF activation due to fragmentation of ovarian tissue (43). We used ovaries from one-week-old females, the stage at which mouse ovaries predominantly contain fully formed PMFs and small population of primary/secondary follicles. This approach also facilitates investigation of direct toxicity of primordial oocytes without the potential confounding effects from damage to growing follicles seen in post-pubertal females. Ovaries were cultured for five days after completion of 48 hours drug treatment, or a total of one week, to ensure observation of long-term primordial oocyte survival rather than delay in apoptosis. To determine doses that significantly deplete primordial oocytes, we exposed ovarian explants from wildtype females to increasing doses of MAFO (0.1-1 μg/ml), CDDP (0.1-0.5 μg/ml), DOX (0.025-0.1 μg/ml) and ETO (0.05-0.5 μg/ml) (Fig. S1). The number of primordial oocytes was reduced compared to vehicle controls following exposure to all tested drugs in a dose dependent manner (Fig. S1).

Significantly reduced survival of primordial oocytes was observed in wildtype ovaries for MAFO at 1 μg/ml (7.3% ± 9.0; P<0.0001), CDDP at 0.5 μg/ml (0.04% ±0.08; P<0.0001) and DOX at 0.1 μg/ml (0.8% ± 0.6; P <0.0001), and to lesser extent for ETO at 0.5 μg/ml (39.0% ±13.7; P = 0.0021) (Fig. 1A,C). In agreement with published data, ETO showed lowest toxicity in juvenile ovaries, suggesting mechanistic differences in toxicity compared to the other topoisomerase II poison tested (DOX). When *Chek2*-/- ovaries were treated with the same doses of drugs no significant reduction in primordial oocyte numbers was observed (Fig. 1A, B, C). We calculated oocyte survival as a percentage of ovarian reserve in untreated wildtype and *Chek2*-/- females. When compared to wildtype controls, survival of primordial oocytes lacking CHEK2 was improved after MAFO (7.3% vs. 107.3%; P = 0.0002), CDDP (0.04% vs. 83.2%; P<0.0001), DOX (2.4% vs. 92.9%; P<0.0001) and ETO (39% vs. 120.7%; P = 0.0006).

**Figure. 1.**
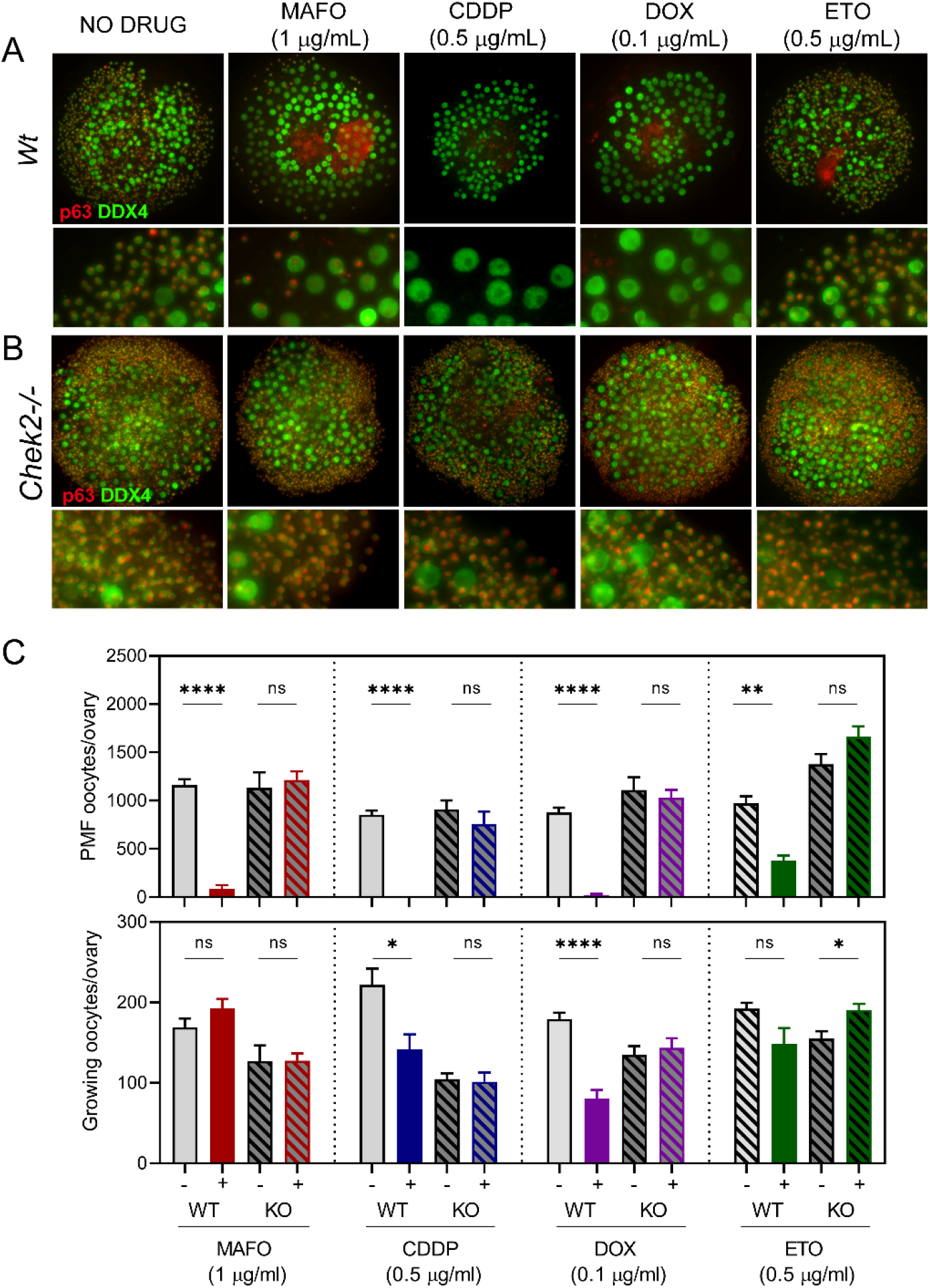
CHEK2 deficiency prevents primordial oocyte depletion after *ex vivo* treatment with MAFO, CDDP, DOX and ETO. Ovaries from wildtype (A) and *Chek2*-/- females after 7-day *ex vivo* culture with MAFO (1 μg/mL), CDDP (0.5 μg/mL), DOX (0.1 μg/mL) or ETO (0.5 μg/mL) were immunostained for oocyte markers DDX4 and p63. Whole ovaries are shown on top and magnified regions below. (C) Primordial and growing oocytes were counted in ovaries harvested after 7-day organ culture. Primordial oocytes are found in PMFs and growing oocytes are found in primary and secondary follicles. Data are expressed as mean ± SEM; *p<0.05, **p<0.01, ***p<0.001, ****p<0.0001 (Mann-Whitney nonparametric test).

Because there have been reports that CDDP and CTX induce hyperactivation of PMFs and their transition to growing follicles (44–46), we counted the larger growing oocytes (diameter > 30 μm), which are typically found in primary and secondary follicles, in 2-week old ovaries treated with each of the four drugs. We observed no significant increase in growing oocyte numbers: to the contrary, in some cases (CDDP, DOX), we observed a reduction in the number of growing oocytes after drug treatments (Fig. 1C, Fig. S1). This indicates that follicle hyperactivation is not the major mechanism of PMF loss in prepubertal ovaries or *ex vivo* explants and is in agreement with other studies (35,47,48). It is possible that MAFO (an analog of CTX) has a different effect on ovaries *ex vivo* than CTX *in vivo*, however both drugs have been shown to generate similar metabolites (37,42). In summary, survival of oocytes in *Chek2* -/- treated ovaries *ex vivo* demonstrate that CHEK2 is directly responsible for elimination of PMFs after treatment with four different chemotherapy drugs and that CHEK2 inhibition is likely sufficient to prevent loss of PMF reserve in cancer patients treated with these drugs.

### CHEK2 deficiency prevents PMF loss after *in vivo* treatment with an acute dose of alkylating chemotherapy drugs

To confirm that abrogation of CHEK2 activity is sufficient to mitigate chemotherapy-induced PMF loss in prepuberal mouse ovaries *in vivo*, one-week-old control (*Chek2*+/-) and *Chek2*-/- females were treated with vehicle or an acute dose of CTX (150 mg/kg) or CDDP (5 mg/kg); both of which are highly ovotoxic (14,41) (49,50). Female weights were recorded before treatment and then weekly until ovaries were collected at week three (two weeks post-treatment) for histological analyses of the PMF reserve. CTX treatment obliterated oocytes in PMF in control females (Fig. 2A,C), while abundant PMFs with oocytes were present in ovaries from CTX treated *Chek2*-/- females (3% vs. 91%; P<0.0001) (Fig. 2A,C). CDDP treatment significantly decreased PMF numbers in ovaries from control animals while abundant PMFs were present in *Chek2*-/- females (37% vs 100%; P = 0.0009) (Fig. 2 B,C). Moreover, improved weight gain was evident in treated *Chek2*-/- pups compared to controls (60% vs. 10%) (Fig. 2D), suggesting that inhibition of CHEK2 activity could potentially alleviate other adverse side effects caused by CTX, although hair loss associated with CTX treatment was not prevented (data not shown).

**Figure 2.**
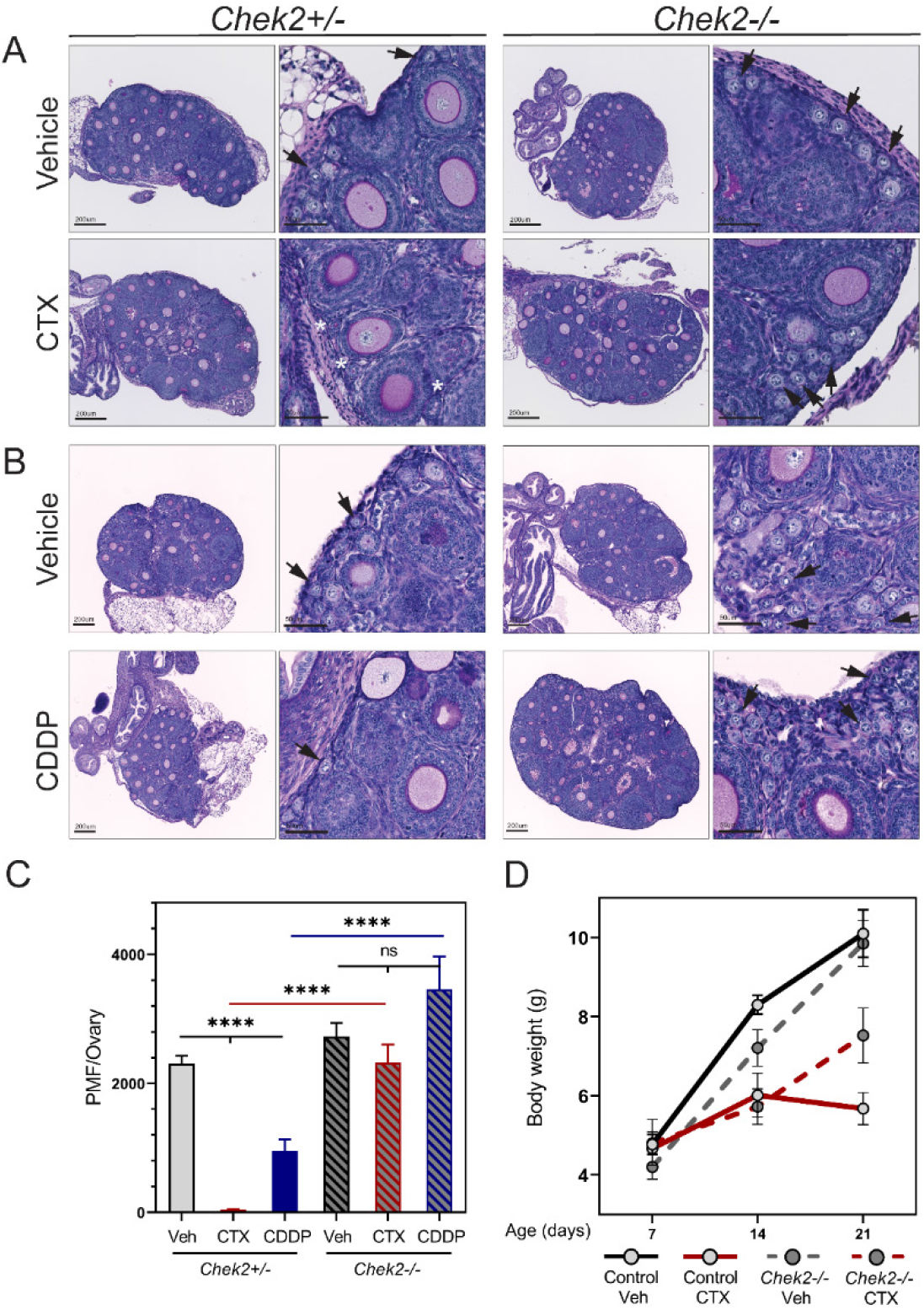
CHEK2 deficient females maintain PMF reserve after *in vivo* treatment with CTX and CDDP. Ovarian histology from *Chek2*+/- and *Chek2*-/- females two weeks after *in vivo* treatment with cyclophosphamide CTX (150 mg/kg) (A) and CDDP (5 mg/kg)(B). Arrows indicate PMFs and asterisks follicle remnants devoid of oocytes. (C) PMF numbers counted in *Chek2*+/- and *Chek2*-/- ovaries 2 weeks post treatment with drugs. (D) Body weight changes in *Chek2*+/- and *Chek2*-/- females treated with CTX. Data are expressed as mean ± SEM; *p<0.05, **p<0.01, ***p<0.001, ****p<0.0001 (Mann-Whitney nonparametric test).

CTX and CDDP treated *Chek2*-/- females retained >90% of their ovarian reserve and showed healthy growing follicles in the ovary at 3 weeks of age, thus were expected to be fertile. In this study we focused on PMF survival rather than fertility outcomes but other studies indicate that oocytes that survive genotoxic treatments can produce healthy offspring (33,34). Taken together, these results indicate that targeting CHEK2 activity *in vivo* is sufficient to protect against CDDP and CTX induced PMF loss.

### CDDP and MAFO treatments cause DNA DSBs in oocytes, which exhibit markers for homologous recombination but not non-homologous end joining repair

CHEK2 has been shown to coordinate elimination of oocytes in response to unrepaired, programmed meiotic and radiation-induced DSBs (26,51). To examine whether the four tested drugs cause DNA DSBs in oocytes, which would activate CHEK2 signaling, vehicle- and drug-treated wildtype ovaries were immunostained for general DNA DSB marker γ-H2AX. After 24 hours of treatment, abundant γ-H2AX foci were present in oocytes treated with MAFO and CDDP but not with DOX or ETO (Fig. 3A, D). Since DSB can be repaired by high fidelity homologous recombination (HR) or more error-prone non-homologous end joining (NHEJ), ovaries were immunostained for HR marker RAD51 and NHEJ marker 53bp1.

**Figure 3.**
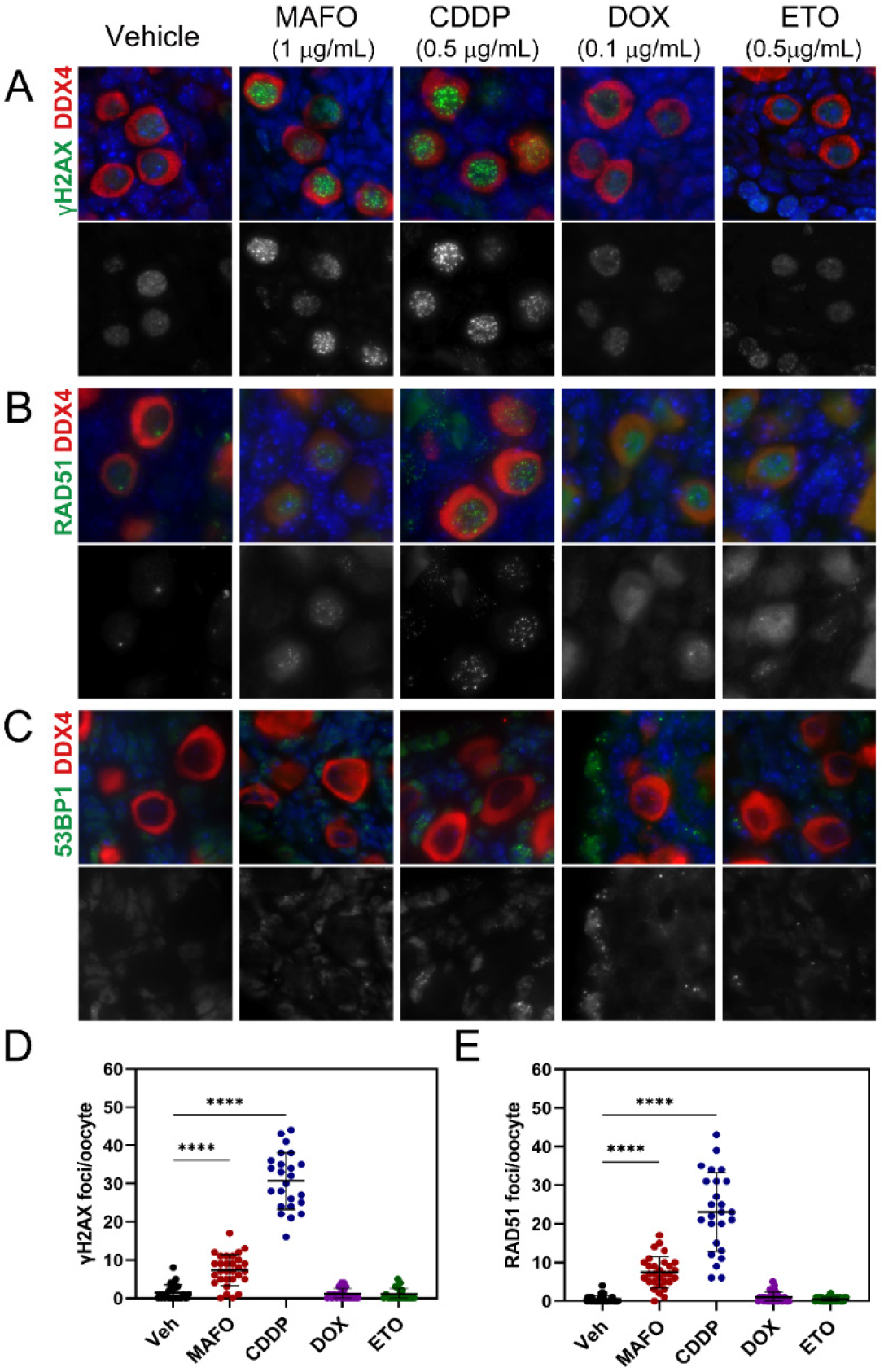
MAFO and CDDP treatment of oocytes causes DNA DSBs that activate HR, but not NHEJ, repair. Ovaries were exposed to chemotherapy drugs *ex vivo* for 24 hrs. Ovarian sections were immunostained for (A) general DNA damage marker γH2AX and DNA repair markers (B) RAD51 for HR and (C) 53BP1 for NHEJ. Quantification of DSBs per oocyte using (D) γH2AX and (C) RAD51 markers. Data are expressed as mean ± SEM; *p<0.05, **p<0.01, ***p<0.001, ****p<0.0001 (one-way ANOVA, Bonferroni multiple comparison test).

RAD51 foci were present in oocytes treated with MAFO or CDDP, indicating that these drugs activate the HR repair pathway (Fig. 3B, E). There were no RAD51 foci in DOX and ETO treated oocytes, consistent with the γ-H2AX staining results (Fig. 3B, E). Oocytes in postnatals ovaries are arrested at dictyate stage of meiotic prophase I, comparable to mitotic G2 phase. In this phase, HR is believed to be the predominant repair pathway due to the presence of sister chromatids, but NHEJ repair is also thought to be active. A previous report shows that DNA-PKcs, a component of NHEJ repair, is detected in only 10% of primordial oocytes after radiation-induced DNA damage (52). Another study reported DNA-PKcs activation in 30-60% of oocytes after CTX treatment (53). To determine the activation of NHEJ in oocytes after chemotherapy, ovaries were immunostained with 53BP1(Fig. 3C) (54,55). 53BP1directs DSB repair towards NHEJ and localizes to DSB sites forming discrete foci that colocalize with γ-H2AX (54–56). Remarkably, there was no evidence for 53BP1 foci in oocytes treated with any of our tested chemotherapy drugs. However, they were readily detected in somatic cells in the ovary (Fig. 3C). This indicates that DNA damage in oocytes is predominantly, if not exclusively, repaired by high fidelity HR, which has lower risk of generating deleterious mutations, as was shown for radiation (52).

Lack of γ-H2AX in oocytes after DOX and ETO treatments suggests that DSBs are not the predominate type of damage activating CHEK2 signaling leading to oocyte apoptosis.

Taken together we show that CDDP and MAFO induce lethal DSBs in oocytes. Repair of these DSBs is initiated by high fidelity HR but cannot be completed due to activation of proapoptotic response coordinated by CHEK2. Moreover, our results confirm previous reports that topoisomerase II poisons such as DOX and ETO do not induce DSBs in oocytes (57), but rather in ovarian somatic cells that are actively proliferating. Nevertheless, primordial oocyte elimination is still observed in CHEK2-proficient ovaries, indicating that DOX and ETO affect primordial oocyte survival by a different mechanism. Taken together our results indicate that inhibition of CHEK2 activity in the chemotherapy-treated ovary prevents elimination of primordial oocytes with and without DNA DSBs by providing more time for DNA repair and oocyte survival.

### Damage induced by CDDP and MAFO does not induce hyperphosphorylation of TAp63, but triggers activation of TRP53-dependent apoptosis

In response to radiation-induced damage, CHEK2 phosphorylates two proapoptotic factors; TAp63 specifically in oocytes and TRP53 in all cell types (26,32,58). Unlike TRP53, which requires phosphorylation to avoid degradation, TAp63 is constitutively expressed in oocytes and maintained in an inactive form until phosphorylated (32,58,59). In response to radiation, TAp63 is phosphorylated by CHEK2 and Casein Kinase 1 delta (CK1δ) leading to a mobility shift visible by western blot. In the absence of CHEK2, TAp63 remains unphosphorylated and inactive (26) (Fig. 4A and Fig. S2C). Previous reports suggest that inactivation of TAp63 is sufficient to protect oocytes from CDDP but not CTX induced damage (33,34). Since MAFO and CDDP induce DSBs and trigger CHEK2-dependent apoptosis, phosphorylation and activation of TAp63 is expected. However, TAp63 shift was absent after 24 hours of MAFO or CDDP treatment, even though activated TRP53 and γH2AX were detected by western blot (Fig. 4A). These results suggest that damage induced by MAFO (2.4 μM) and CDDP (1.6 μM) does not lead to the rapid phosphorylation of TAp63 observed after radiation-induced damage. However, TAp63 phosphorylation has been reported in oocytes after treatments with a higher dose of CDDP (10 μM) (32) or after CTX treatment *in vivo* (53). Therefore, it is possible that DNA damage caused by alkylating agents triggers activation of TAp63 with different dynamics or by a different mechanism. To determine whether TAp63 is involved in MAFO- and CDDP-induced primordial oocyte elimination downstream of CHEK2, we utilized a mouse model with a mutation in the *Trp63* gene (Fig. S2A). Here, serine 621 was replaced with alanine (S582 in human) at the TAp63 C-terminus, which was previously showed to be phosphorylated by CHEK2 (26,32). This mutation prevents activation by CHEK2 but does not affect TAp63 expression (Fig. S2C) and results in increased resistance of oocytes to low dose of radiation (0.5Gy) (Fig. S2B). *TAp63A/A* ovaries were exposed *ex vivo* to MAFO and CDDP at the same doses and regimen as wildtype and *Chek2*-/- ovaries (Fig. 4B, C and Fig. 1). Primordial oocyte numbers were reduced in treated *TAp63A/A* ovaries and when compared to *Chek2*-/-: only ~21.9% survived (vs. ~107.3% P <0.0001) after MAFO and ~12.7% (vs. ~83.3% P <0.0001) after CDDP. The reduced number of surviving primordial oocytes in *TAp63A/A* mutants suggests that other CHEK2 targets contribute to primordial oocyte elimination after CDDP- and MAFO-induced damage. CHEK2 is known to phosphorylate TRP53 in response to DNA DSBs, and TRP53 expression was detected in ovaries treated with radiation, MAFO, and CDDP (Fig. 4A). Interestingly, TRP53 was still detected by western blot in irradiated ovaries in *Chek2*-/- mice, although at lower levels than in the wildtype (Fig. S2C), indicating that CHEK1 or other kinases can still activate TRP53 in the absence of CHEK2. However, it remains unclear whether the remaining TRP53 is activated in oocytes or ovarian somatic cells. To test if TRP53 contributes to primordial oocyte elimination after MAFO and CDDP-induced damage, oocyte survival was tested in double mutant ovaries lacking activity of both TAp63 and TRP53. Compared to *Trp63A/A* single mutant, primordial oocyte survival was significantly improved in *Trp63A/A Trp53*-/- double mutant ovaries (21.9% vs. 84.4% P=0.002 after MAFO and 12.7% vs. 99.4% P=0.0013 after CDDP) (Fig. 4C). This confirms that TRP53 is activated in the ovaries after MAFO and CDDP-induced damage and that it contributes to primordial oocyte elimination. Because CHEK2 is expressed in all ovarian cells, further studies are required to determine if TRP53 activity, critical to primordial oocyte elimination, comes from surrounding ovarian somatic cells or the oocyte itself. Taken together, genetic analysis reveals the critical role for CHEK2 in coordinating PMF loss after MAFO and CDDP-induced damage by activating two proapoptotic factors TAp63 and TRP53.

**Figure 4.**
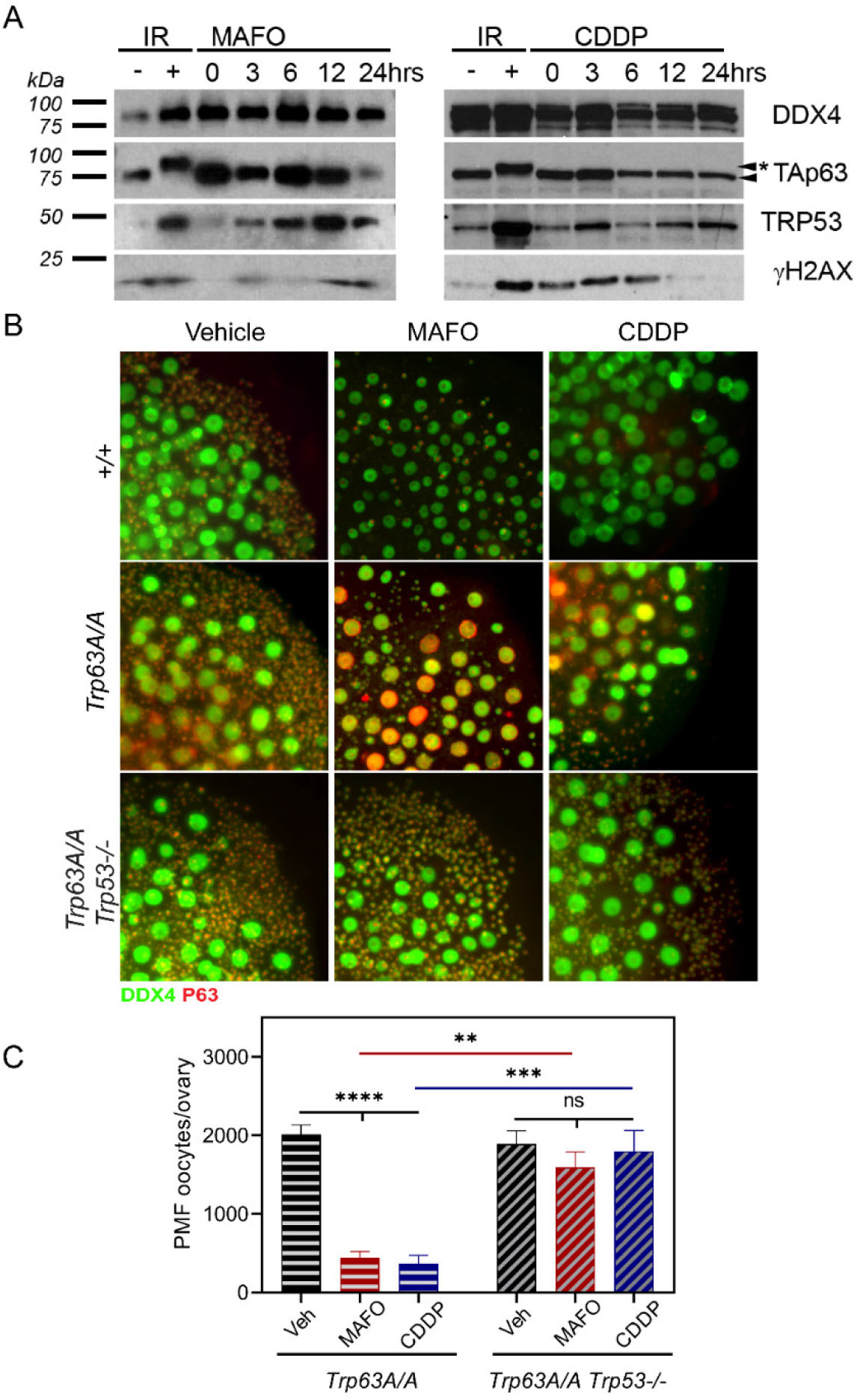
Inhibition of TAp63 phosphorylation by CHEK2 is not sufficient to prevent complete primordial oocyte elimination in response to MAFO and CDDP treatments indicating a role for TRP53. (A) Protein extracts were collected from MAFO and CDDP-treated ovaries at different time points and analyzed by western blot (MAFO at 1μg/ml at CDDP at 0.5μg/ml). TAp63 phosphorylation leads to mobility shift observed 6 hrs after radiation (IR 0.5Gy). In contrast no shift is observed in CDDP and MAFO treated ovaries up to 24 hrs post treatment. Increased levels of TRP53, indicative of its phosphorylation and stabilization, are observed as early as 3 hrs post drug treatments. DDX4 is an oocyte marker and γH2AX is a DNA damage marker. (B) Primordial oocytes expressing TAp63 carrying a mutation at its CHEK2 phosphorylation site (*Trp63A/A*) are not fully resistant to MAFO and CDDP toxicity while double mutant *Trp63A/A Trp53*-/- oocytes lacking both CHEK2 phosphorylation targets show high resistance and almost normal survival. (C) Primordial (PMF) oocytes were counted in ovaries harvested after 7-day organ culture. Data are expressed as mean±SEM; ****p<0.0001 (one-way ANOVA, Kruskal–Wallis for nonparametric data).

### Transient pharmacological inhibition of CHEK2 improves primordial oocyte survival after CDDP and MAFO treatment

Genetic ablation of CHEK2 activity prevents depletion of PMF reserve in mice treated with chemotherapy drugs (this study) and radiation (26). Moreover, pharmacological inhibition with CHEK2 Inhibitor II (BML-277) reduced oocyte loss after radiation and resulted in healthy offspring from surviving oocytes (36). Because of similarities between CHEK1 and CHEK2, most available inhibitors block the activity of both kinases, although with different affinities. CHEK1 is an essential kinase which coordinates DDR and cell cycle progression in all dividing cells (60). CHEK1/2 inhibitors have been shown to potentiate the effects of genotoxic chemotherapy drugs against cancer (61–65) and some are being tested in clinical trials. Therefore, inhibitors blocking CHEK2 and CHEK1 could have the added benefit of improved cancer cell elimination in addition to oocyte protection. To test the ability of other CHEK1/2 inhibitors to prevent oocyte depletion following radiation and chemotherapy treatments, four inhibitors shown or predicted to target CHEK2 were tested; AZD7762 (IC50 CHEK2 10nM and CHEK1 5nM), CCT241533 (IC50 CHEK2 3nM and CHEK1 245nM), LY2606368 (IC50 CHEK2 8nM and CHEK1 <1 nM), and PF477736 (IC50 CHEK2 47 nM(Ki) and CHEK1 0.49 nM(Ki); IC50 provided by suppliers). The pharmacokinetics of these inhibitors in mouse are unknown. Therefore, we used our *ex vivo* ovary culture system to better control the timing and dosing of drugs and inhibitors without the confounding influence of transport and metabolism found *in vivo*. First, ovaries were treated with increasing doses of inhibitors in combination with ionizing radiation (0.5 Gy) (Fig. 5A). Ovaries were cultured with inhibitors for 2 hours prior to treatment to assure cellular presence of the inhibitor before DNA damage is induced and for additional 48 hours after radiation. Oocyte numbers were counted in immunostained whole ovaries after an additional 5 days of culture without any inhibitors to ensure quantification of oocyte survival and not a delay in apoptosis. Treatment with CCT241533 (0.05, 0.5 and 5μM), LY2606368 (0.1, 1 and 10μM), and PF477736 (0.5, 5 and 50μM) had no significant impact on primordial oocyte survival after radiation at any dose tested (Fig. 5A) indicating failure to sufficiently block CHEK2 activity in oocytes. AZD7762 treatment improved primordial oocyte survival compared to vehicle-treated irradiated ovaries (Fig. 5A). We calculated survival as a percentage of oocyte reserve in non-irradiated ovaries. Survival was estimated at 57.3% after 10μM AZD7762 (P=0.001) vs. 10.1% with vehicle. Survival at lower doses of inhibitor (0.1 and 1μM) was improved but more variable. Despite increased primordial oocyte survival, ovarian explant growth was impaired in AZD7762 treated ovaries. Immunostaining for DNA damage marker γH2AX and TUNEL for apoptosis shows that AZD7762 treatment alone causes accumulation of DNA damage and increased apoptosis in ovarian somatic cells which was further enhanced by radiation (Fig. 5B). Since *Chek2*-/- ovaries show normal growth after radiation, these results suggest that AZD7762 inhibited CHEK1 kinase which interfered with cell cycle progression in proliferating stromal and granulosa cells. This led to accumulation of DNA damage and triggered apoptosis in somatic cells but not in oocytes.

**Figure 5.**
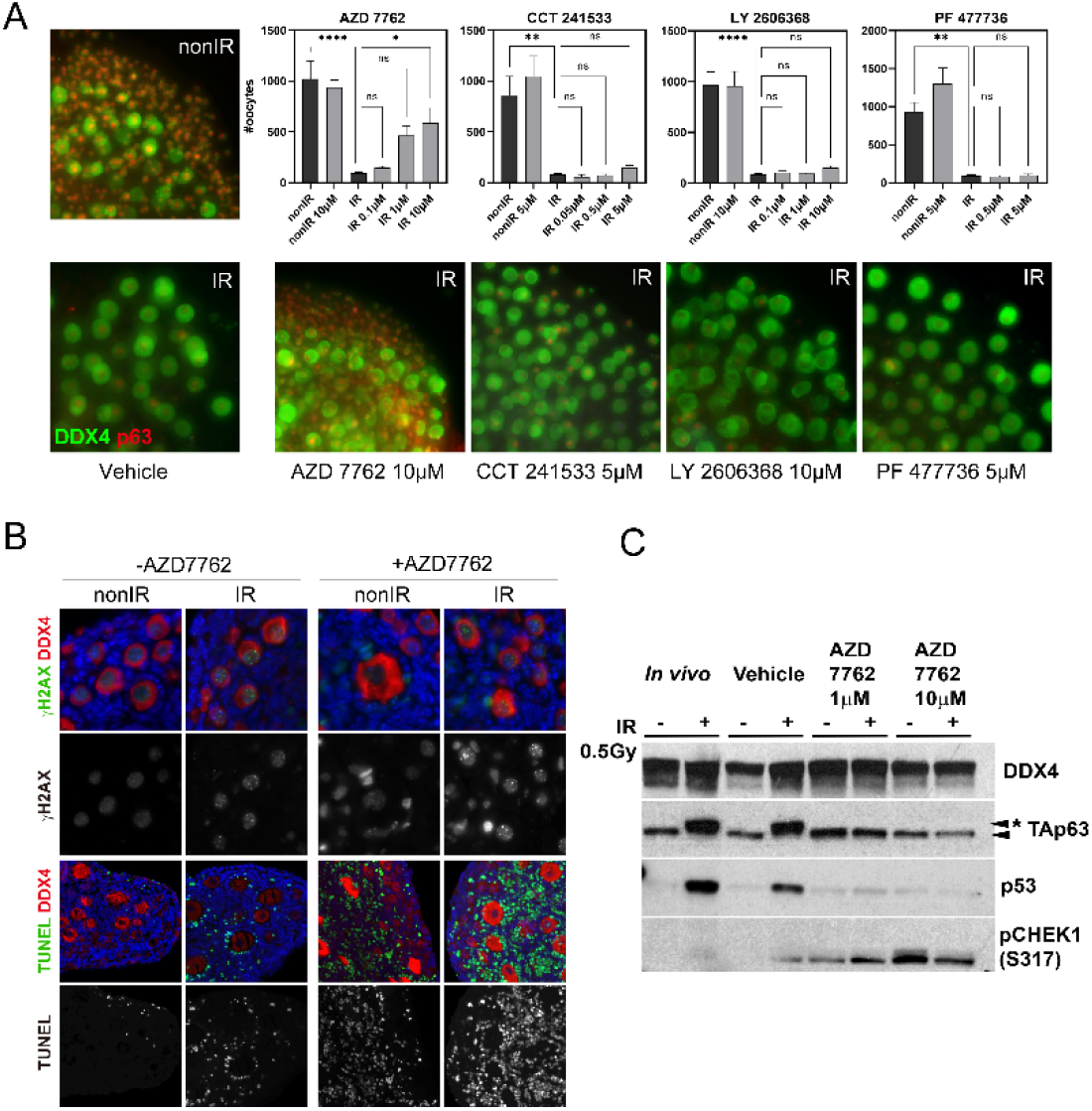
CHEK1/2 inhibitor AZD7762 reduces primordial oocyte loss in prepubertal ovaries after radiation (IR) but exhibits cytotoxic effects in proliferating somatic cells. (A) Ovaries were treated with CHEK2 inhibitors *ex vivo* for 2 hrs prior to IR and for 24 hrs post IR. After 24 hrs inhibitors were withdrawn and ovaries cultured for 6 days. AZD7762 but not CCT241533, LY2606368 or PF477736 showed protection of PMF oocytes after 7-day culture. Graphs show numbers of oocytes per ovary present after treatment with increasing doses of inhibitors. Data are expressed as mean ± SEM; *p<0.05, **p<0.01, ***p<0.001, ****p<0.0001 (one-way ANOVA, Kruskal–Wallis with Dunn’s multiple comparison test for nonparametric data). Below, panels show examples of treated ovaries immunostained with oocyte markers DDX4 and p63. (B) AZD7762 treatment causes increased levels of γH2AX (green) staining in the absence of IR. TUNEL staining (green) for apoptotic cells shows increased apoptosis in samples treated with AZD7762. Oocyte marker DDX4 (red). (C) Ovarian protein extracts were collected 6 hrs after IR with 0.5Gy with and without AZD7762 treatment and were analyzed by western blot. TAp63 mobility shift (asterisk) indicative of phosphorylation and p53 expression are not detected in inhibitor treated ovaries. Increased levels of pCHEK1(S317) are present only in AZD7762 treated ovaries.

To validate that AZD7762 prevents primordial oocyte death by inhibition of CHEK2-dependent signaling, we performed western blots of TAp63 and TRP53 as before (66,67). TAp63 was phosphorylated after radiation which resulted in the expected mobility shift and TRP53 was readily detected (Fig. 5C). In ovaries treated with AZD7762, TAp63 remained unphosphorylated and TRP53 was reduced which confirms that AZD7762 efficiently inhibited CHEK2 activity in oocytes and prevented TAp63-activation. CHEK1 is phosphorylated on Serine 317 by ATR kinase in response to DNA damage (60), which we were able to detect in all AZD7762-treated cells (Fig. 5B). To test whether AZD7762 could also protect primordial oocytes against chemotherapy drugs, ovaries were co-treated with AZD7762 (1μM and 10μM) and MAFO or CDDP (Fig. 6A, B). Oocyte survival was calculated as a percentage of ovarian reserve in untreated ovaries. Oocyte survival increased in 1μM AZD7762-treated ovaries compared to vehicle control following co-treatment with MAFO (59.6 vs. 4.5% P=0.0003) and CDDP (63.1 vs. 3.9% P=0.0006). Despite improved oocyte survival, retardation of ovarian growth was also observed in co-treatment with chemotherapy drugs (Fig. 6A). This suggests that despite its ovoprotective and cancer sensitizing properties, AZD7762 causes additional cytotoxic side effects in the ovary. AZD7762 cardiotoxicity was reported in clinical trials which led to their termination (68–72). In summary, these data show that transient inhibition of CHEK2 is a feasible approach to protect ovarian PMF reserve during genotoxic chemotherapy treatments providing that a highly specific CHEK2 inhibitor is used.

**Figure 6.**
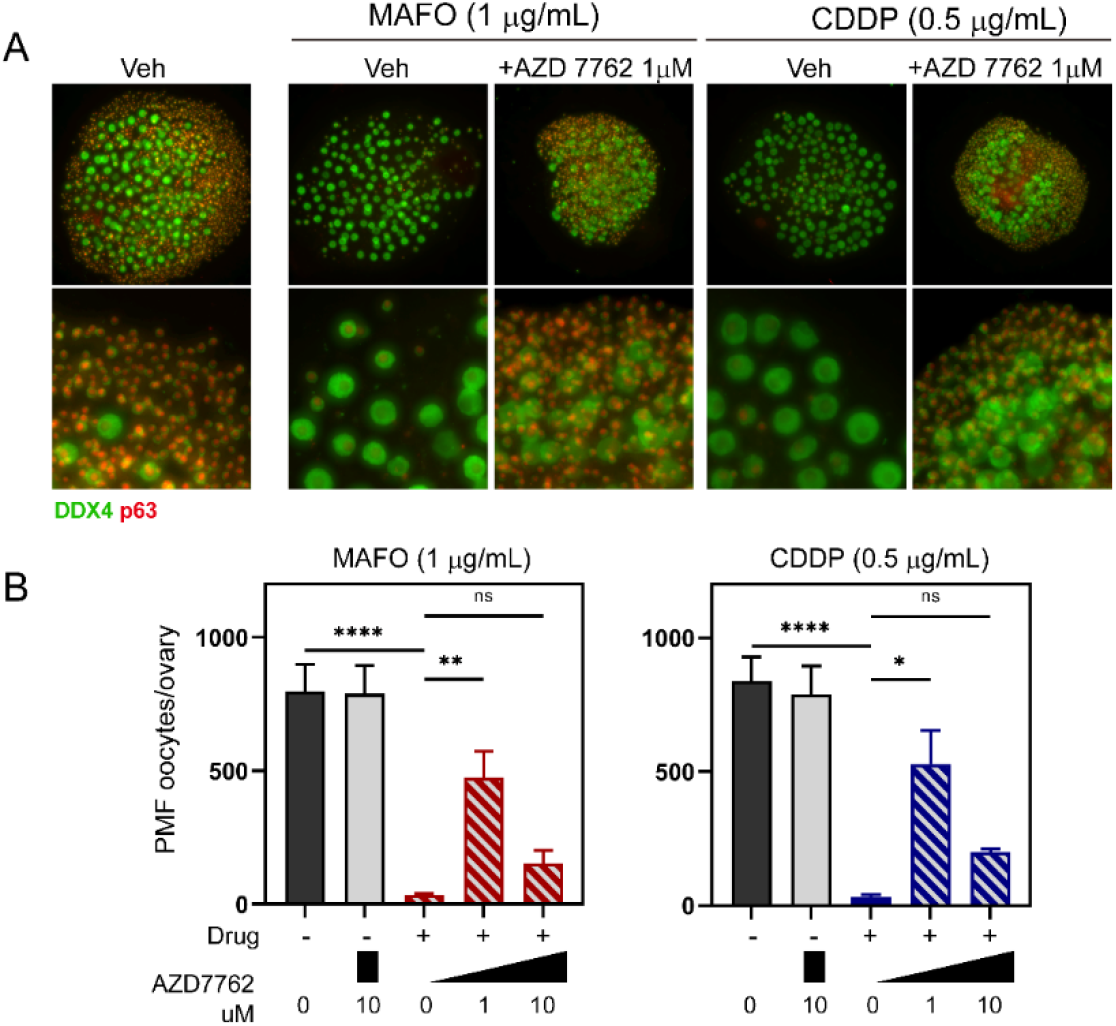
Co-treatment with AZD7762 reduces primordial oocyte loss in prepubertal ovaries after treatment with chemotherapy drugs. (A) Example ovaries co-treated with AZD7762 and MAFO or CDDP in *ex vivo* organ culture. AZD7762 showed protective effect and presence of primordial oocytes after 7-day culture. Oocyte markers DDX4 (green) and p63 (red). (B) Graphs show numbers of primordial oocytes per ovary present after co-treatment with increasing doses of AZD7762 inhibitor. Data are expressed as mean ± SEM; *p<0.05, **p<0.01, ***p<0.001, ****p<0.0001 (one-way ANOVA, Kruskal– Wallis with Dunn’s multiple comparison test for nonparametric data).

## DISCUSSION

Genotoxic cancer treatments can damage and diminish PMF reserve in cancer patients causing POI and fertility problems, posing challenges that reveal the limitations of our understanding of the basic biology of oocyte in the context of follicle environment. Mouse studies demonstrate that PMF loss can be prevented in females treated with genotoxic agents, and that these mice remain fertile and produce healthy pups. However, it remains unclear what might be the safest and most effective way to mitigate PMF loss in female cancer patients. In this study, we show that CHEK2 kinase is responsible for coordinating elimination of primordial oocytes after damage with several chemotherapy drugs, making it an attractive target for the development of ovoprotective treatments. Genetic ablation of CHEK2 protected >90% of PMF reserve in females treated with two very ovotoxic drugs, CTX and CDDP, demonstrating that blocking CHEK2 is sufficient to protect PMF reserve in mice. Although fertility was not assessed in this study, evidence from prior work and other studies indicate that primordial oocytes that survive genotoxic insults do indeed support normal ovarian function and fertility (26,33,35,36). Moreover, CHEK2 inactivation also almost completely protected primordial oocytes from MAFO, DOX and ETO toxicity in *ex vivo* organ culture system. Although DOX and ETO are not as ovotoxic as alkylating agents in human (38,73,74), they are used in many combination therapies (75–77). Thus, these findings suggest the protective effect of CHEK2 inhibition is likely to be beneficial for a broad spectrum of patient treatments. Further, we demonstrate that a CHEK1/2 inhibitor previously shown to sensitize cancer cells to chemotherapy significantly improved primordial oocyte survival after either radiation or treatment with alkylating agents. Finally, our genetic dissection of the signaling mechanisms responsible for PMF elimination post genotoxic insult establishes CHEK2 as a master regulator of this process and reveals parallel and redundant downstream pathways that involve both TAp63 and TRP53.

How different chemotherapy drugs inflict damage in ovaries and oocytes specifically, and how this damage triggers primordial but not growing oocyte elimination is still not fully understood. Chemotherapy drugs can damage DNA directly and indirectly by increasing oxidative stress. Moreover, oxidative stress causes other cellular damage via protein and lipid peroxidation, which can also lead to cell death (78). Here, we show that alkylating agents such as CDDP, MAFO, and CTX induce abundant DNA DSBs (as previously reported (49,79)) in oocytes that in turn trigger CHEK2 activation and subsequent loss of PMFs by oocyte apoptosis. In contrast, we did not detect a significant increase in DSB markers in primordial oocytes after DOX or ETO treatment, suggesting a different mode of toxicity for these agents. It is possible that DOX induces late DNA damage in oocytes indirectly by increasing oxidative stress and reactive oxygen species (ROS) (80–83). Indeed, co-administration of drugs reducing oxidative stress have been shown to decrease its ovotoxicity (84–86) and γH2AX-positive oocytes were reported after *in vivo* treatment with DOX (57,87). The improved survival of primordial oocytes in *Chek2*-/- ovaries after DOX and ETO exposure suggests that these drugs induce CHEK2 signaling independently of nuclear DNA damage, most likely through increased oxidative stress (81,88–91). Studies show that DOX and ETO treatment are most damaging to actively proliferating granulosa cells (38,40,41,57). It remains unclear whether loss of PMFs after DOX and ETO is driven by the response from the oocyte or surrounding granulosa cells and if damage to one cell type is communicated to the other within the follicle (discussed below). Because CHEK2 is expressed in all cells in the ovary, its inactivation in oocytes, granulosa cells or both may be responsible for observed primordial oocyte survival.

The primordial oocyte response to DNA DSBs induced by radiation involves CHEK2 and its downstream target TAp63, with TRP53 shown to be dispensable (26,92,93). Both genetic and biochemical evidence indicate that phosphorylation by CHEK2, followed by CK1δ, are needed to fully activate TAp63 (26,32). Hyper-phosphorylated TAp63 can then tetramerize and function as a transcription factor (32,58,59,94). However, the role of CHEK2 in the response to alkylating agents has been unclear due to discrepancies in TAp63 hyper-phosphorylation status and its requirement for oocyte elimination. TAp63 hyper-phosphorylation is not evident after CDDP and MAFO treatments (this study and (34,95)) but has been reported after *in vivo* administration of CTX (53), whereas TAp63 null oocytes are resistant to CDDP but not to CTX (33,34). A study using an array of DDR inhibitors against ATR, ATM, CHEK1 and CHEK2 kinases suggested that radiation and CDDP activate TAp63 by two distinct signaling cascades; ATM→CHEK2→TAp63 and ATR→CHEK1→TAp63, respectively (34). Since many DDR inhibitors show activity towards other kinases (96,97) and there is a cross-talk between ATM and ATR pathways (98–100) it is possible that survival attributed to ATR→CHEK1 inhibition is due to off-target inhibition of CHEK2. In this study, we observed that the inhibitor AZD7762 targeted both CHEK1 and CHEK2 kinases while other predicted CHEK2 inhibitors failed to inactivate CHEK2 and showed no protective effects. While inhibitor studies are helpful in initial identification of signaling cascades involved in cellular responses, genetic studies are necessary to determine precise mechanistic roles for individual proteins (97), for example, as shown for imatinib and ABL1 kinase. Initial studies with the inhibitor imatinib provided conflicting results about its protective effects against CDDP while genetic study demonstrated that ABL1 kinase is dispensable for elimination of CDDP-damaged oocytes (40,95,101–103). Here, using a genetic approach we demonstrate that CHEK2 is directly responsible for elimination of primordial oocytes damaged by both CDDP and CTX and that CHEK1 activity does not significantly contribute to oocyte elimination even in the absence of CHEK2.

Since previous reports suggested *TAp63*-/- oocytes are resistant to CDDP but not to CTX (33,34), and that *Chek2*-/- oocytes are resistant to both (this study), other types of cellular damage and/or other targets of CHEK2 must be involved in signaling pathways leading to elimination of oocytes damaged by CTX. At least two putative mechanisms underlying its ovotoxic effect have been proposed: 1) accelerated activation of PMFs causing so called follicle “burn-out” via the PTEN/AKT/FOXO3a pathway (44–46,50); and 2) direct damage to the oocyte triggering apoptosis via the DDR pathway (35,48,49). However, the relative contribution of these two mechanisms to PMF loss remains controversial (35,44–46,48,104,105). CTX treatment in pre- and post-pubertal mice demonstrated that primordial oocytes are lost within 3 days without an increase in the number of growing follicles, indicating that follicle burn-out is not the major mechanism regardless of female age (35). This study and others show that CTX and its derivative MAFO are potent inducers of DSBs and apoptosis in oocytes (49,106).

Therefore, primordial oocyte survival in ovaries lacking CHEK2—the essential DSB response kinase—after both MAFO (*ex vivo*) and CTX (*in vivo*) treatments strongly suggests that DNA damage is the major mechanism underlying primordial oocyte loss after CTX treatment. Even though radiation, CDDP, and MAFO/CTX all induce DSBs in oocytes, the mechanisms for the ensuing activation of TAp63 and TRP53 are different. We show that despite inducing slightly lower levels of DSBs than CDDP, MAFO treatment specifically activates TRP53-dependent apoptosis suggesting that DSB-independent cellular damage induces TRP53. Full survival of primordial oocytes in MAFO treated ovaries was achieved only after both TRP53 and TAp63 were deleted, suggesting that MAFO-induced damage activates both targets of CHEK2 independently or that they can substitute for each other. Oxidative stress could play a role as CTX and its derivatives produce reactive metabolites such as phosphoramide mustard and acrolein that cause overproduction of ROS which is known to activate CHEK2→TRP53 pathway (37,88,89,107,108). CDDP and radiation also induce oxidative stress (109–111), yet they do not activate TRP53-dependent primordial oocyte loss even in the absence of TAp63 (33,34). In contrast to oocyte-specific expression of TAp63, TRP53 is ubiquitously expressed in the ovary as is CHEK2. Therefore, damage to granulosa cells in the PMF or other cell types in the ovary could potentially trigger oocyte loss through intercellular signaling independent of TAp63 activity in the oocyte, which is prevented by CHEK2 or TRP53 inactivation.

Surprisingly, in contrast to *TAp63*-/- null genetic model, we observed that primordial oocytes expressing the phosphomutant TAp63A/A—lacking CHEK2 phosphorylation site associated with hyper-phosphorylation— were still largely sensitive to CDDP even though CHEK2-deficient oocytes were fully resistant to the same dose. This difference in oocyte survival in the mutants raises two intriguing possibilities; 1) another modification downstream of CHEK2 contributes to TAp63 activation without inducing a hyper-phosphorylated state (as alluded by (34)) or 2) other proapoptotic proteins regulated by CHEK2 are involved. Because TAp63 substitutes for TRP53 function during elimination of oocytes with unrepaired meiotic DSBs (26), we favor the latter explanation. Indeed, we show that deleting *Trp53* in TAp63 phosphomutant background results in CDDP resistance, indicating that TRP53 can contribute to elimination of CDDP-damaged primordial oocytes; however it remains unclear how. It is possible that TAp63 phosphomutant protein triggers oocyte apoptosis by forming an active heterotetramer with TRP53. Although heterotypic interactions have not been detected between wildtype TRP53 and TAp63 in somatic cells (112–114), further investigations are needed to exclude this possibility in oocytes.

Phenotypic analyses of selective mouse mutants reveal a complex network of signaling, with responses depending on the type of drug and potentially type of the cellular damage (Fig. 7A, B). *Chek2*-/- oocytes survive radiation, CDDP, CTX, DOX, and ETO treatments. Downstream of CHEK2, TRP53 is involved in elimination of oocytes exposed to CTX but not CDDP or radiation, while TAp63 seems to be solely responsible for elimination of oocytes exposed to low dose radiation and CDDP (Fig. 7C). TRP53 and TAp63 are known to induce expression of multiple target genes involved in apoptosis, including *Bbc3* (*Puma*) and *Pmaip1* (*Noxa*) (25,115–117). Oocytes lacking PUMA are fully resistant to CTX and CDDP (33), but only moderately resistant to radiation (25). To fully protect oocytes from radiation-induced damage, both PUMA and NOXA need to be inactivated (25). FOXO3a has been suggested to directly activate *Puma* expression in CTX-treated ovaries, thus explaining TAp63-independent oocyte apoptosis (33,118–120). Importantly, FOXO3a was shown to be essential for DNA damage-induced apoptosis dependent on ATM, CHEK2 and TRP53 activity in somatic cells (121,122). Therefore, it is plausible that CTX/MAFO induced damage activates TRP53 via FOXO3a→CHEK2 signaling. FOXO3a is also involved in cellular response to oxidative stress (123–125) thus increased production of ROS in CTX-treated cells can lead to activation of FOXO3a→TRP53→PUMA signaling (Fig. 7A). While these differences in downstream signaling likely reflect nuances in types and/or levels of cellular damage induced by various chemotherapies, CHEK2 nonetheless emerges as a master regulator of primordial oocyte survival or death after exposure to all tested genotoxic agents.

**Figure 7.**
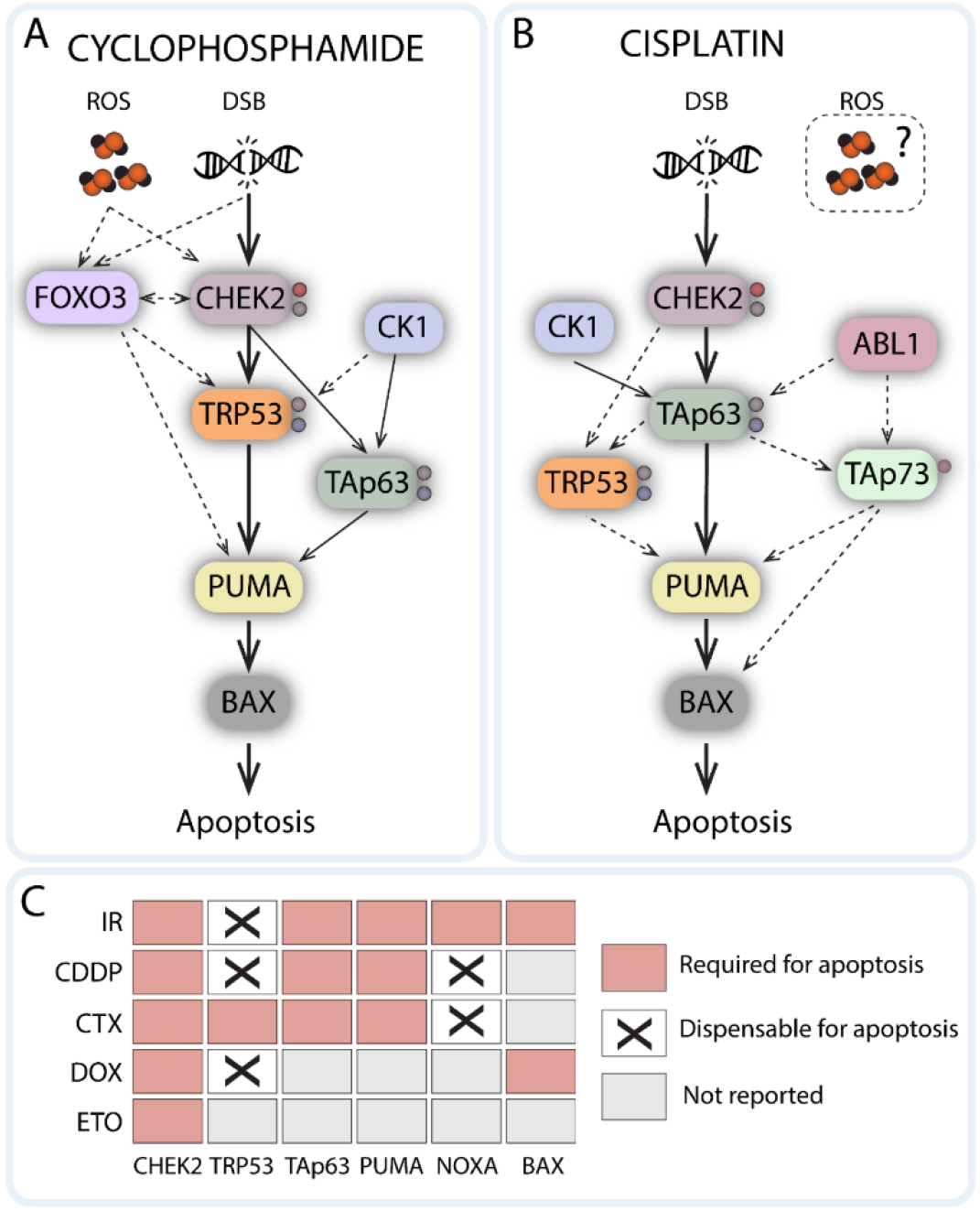
Schematic model of DNA damage signaling in primordial oocytes after cyclophosphamide (CTX) and cisplatin (CDDP) exposure. Solid arrows indicate validated interactions, dashed arrows indicate proposed interactions. DSB: Double Strand Breaks; ROS: Reactive Oxygen Species. C) Matrix illustrating the role of individual proteins in oocyte apoptosis in response to IR and chemotherapy drugs revealed by mutant phenotypes. DOX: Doxorubicin, ETO: Etoposide

In contrast to oocyte specific activity of TAp63, CHEK2 and TRP53 are both also expressed in ovarian cells and thus expected to be involved in DDR response in the granulosa cells as well as in oocytes. Within the follicles, the oocyte and granulosa cells are connected by gap junctions facilitating the bidirectional communication between them (126). These connections ensure exchange of critical metabolites and signaling molecules that support follicular development and survival of both the oocyte and granulosa cells (127–129). While understanding of the molecular mechanisms of radiation and chemotherapy-induced oocyte loss has increased in recent years, the role of intercellular germ cell-soma communication is still emerging. The “bystander effect” where non-targeted cells are affected by adjacent damaged cells has been described for radiation and chemotherapy drugs (130–133), therefore it is possible that granulosa cells with cellular damage affect primordial oocyte survival and *vice versa* through bidirectional communication pathways. Interestingly, gap junctions have been reported to relay DNA damage response to radiation and cisplatin between neighboring cells (134–136) involving ATM, TRP53 and ROS signaling (137–141). Although, it is unclear if cross-communication of DNA damage takes place within the PMF, it has been reported for cumulus-cell enclosed oocytes (142). Further studies, both of clinical responses to chemotherapy and the basic biology of intra-follicle cell communication are required to fully resolve the mechanisms of chemotherapy induced cellular damage and how or where it activates the CHEK2 and TRP53-dependent response.

DNA damage inflicted in oocytes, either directly or indirectly, seems to be the major trigger of PMF elimination and resulting POI after radiation and chemotherapy; therefore, blocking the DDR signaling responsible for this elimination is a promising strategy to prevent POI. Mouse genetic studies have revealed very few targets in the DDR pathway that prevent or minimize elimination of primordial oocytes. Inactivation of proapoptotic factors PUMA and NOXA effectively protected primordial oocytes from radiation and chemotherapy induced damage (25,33,49). However, blocking apoptosis directly may pose an issue for cancer treatment and development of resistance to therapy (143). Inhibition of TAp63 and TRP53, protects primordial oocytes from low dose radiation and selected chemotherapies (25,33,92,93) but targeting transcription factors remains challenging and there are currently no specific inhibitors for TAp63 (144). Consequently, inhibition of DDR kinases responsible for triggering oocyte elimination may be the best approach. CHEK2 and CK1δ are needed for activation of TAp63 in response to DSBs and are both also involved in regulation of TRP53 (67,145–147). Inhibition of their activity genetically or pharmacologically protects primordial oocytes from radiation and selected chemotherapies (26,32). There are six members of the casein kinase family involved in diverse cellular processes including tumorigenesis (148–150) therefore CK1δ specific inhibitors (151) would need to be developed to avoid off-target effects. Using mouse models and CHEK2 inhibitors, we show here that specific inhibition of CHEK2 could be an effective method to prevent chemotherapy-induced loss of ovarian follicle reserve. Moreover, specific inhibition of CHEK2 may also improve cancer treatment outcomes, as DDR kinase inhibitors can sensitize cancer cells to chemotherapy (152). *Chek2*-/- mice are viable, fertile, and do not develop tumors. This suggests that transient inhibition of CHEK2 would be a safe strategy (66,147). While germline mutations in *CHEK2* have been found in patients with breast, colon, and prostate cancers, these appear insufficient to drive cancer progression in the absence of the other cancer susceptibility genes (153). Clinical interpretation of germline *CHEK2* variants is variable, as studies reported their association with increased, decreased, or no risk in various cancers (153). Therefore, further studies are needed to assess the efficacy and safety of transient CHEK2 inhibition in animal models with genetic predisposition to cancer. Comparative analyses of the prevalence of POI in female cancer patients carrying deleterious mutations in *CHEK2* vs. control cohort may also provide insight into the role of CHEK2 in human oocytes’ sensitivity to genotoxic agents and relevance to chemotherapy induced POI as a predictive marker. Recent GWAS studies identified CHEK2 as a major factor associated with ovarian aging and the onset of natural menopause in women (154,155). Animal model studies showed that this is due to a larger PMF reserve found in CHEK2-deficient mice (26,155,156). Furthermore, we showed that CHEK2-deficient primordial oocytes are more resistant to DNA damage induced by radiation (26) and chemotherapy drugs (this study), and we expect that they will be resistant to other types of DNA damaging exposures. Thus, better understanding of how oocytes respond to DNA or other types of cellular damage will contribute to our knowledge of ovarian aging and the maintenance of follicle reserve in women during lifetime exposure to various factors.

## Acknowledgments

This study was supported in part by grant from the V Foundation for Cancer Research to EBF (V Scholar Award V2017-019). The authors would like to thank R. Munroe and C. Abratte of Cornell’s Transgenic Facility for generating the edited mice (supported by a contract CO29155 from the NY State Stem Cell Program (NYSTEM) and R01 GM45415).

## MATERIALS AND METHODS

### Animals

All procedures used in this study were approved by the IACUC at The Jackson Laboratory. C57BL/6J (JAX stock#000664), Tg(Pou5f1-EGFP)2Mnn/J (JAX stock#004654) and *Trp53^tm1Tyj/J^* (JAX stock#002101) were obtained from The Jackson Laboratory. *Chek2^tm1b(EUCOMM)Hmgu^* mice were obtained from KOMP program at JAX. *Trp63*^S621A^ mutant line was generated using CRISPR/Cas9 mediated genome editing as described in Fig. S2. For radiation experiments, 7-day-old females were irradiated using a Cesium-137 gamma irradiator. They were exposed to a total dose of 0.5Gy or 3Gy administered at a rate of (170Rad/min). Ovaries were collected for protein extracts for western blot analysis or fixed in 4% PFA for immunostaining. For chemotherapy drug treatments, 7-day-old females received a single i.p. injection of saline, CDDP (5 mg/kg) (Millipore Sigma) or CTX (150 mg/kg) (Millipore Sigma). Doses were chosen based on previously published studies (49,50). Mice weights were recorded one and two weeks after injection. Ovaries were collected two weeks after injection, fixed in Bouin’s solution for histological analyses.

### Follicle quantification

Bouin’s fixed ovaries were embedded in paraffin and cut into 5 μm serial sections. Sections were stained with Periodic acid-Schiff (PAS) and follicle number was evaluated in every 5th section. Follicle stages were determined using standard methods. Briefly, primordial follicles were surrounded by a layer of squamous pre-granulosa cells; primary follicles had a single layer of cuboidal granulosa cells; secondary follicles had more than one granulosa cell layer but lacked antrum; antral follicles had visible antrum. Final follicle count is represented as the sum of follicles in every 5^th^ section multiplied by a correction factor 5 (157,158). Percentage survival was calculated by dividing the mean of the treatment group counts by the mean of the control group counts and multiplying by a factor of 100.

### Organ culture and drug *ex vivo* treatments

Ovaries collected from 7-day-old pups were cultured on polycarbonate membrane (Whatman Nucleopore Polycarbonate membrane) in MEM (Gibco) supplemented with 10% Fetal Bovine Serum (FBS, Seradigm), 25 mM HEPES (Lonza), and 1X Pen/Strep (Gibco). Ovarian explants were incubated at 37°C, 5% CO_2_, and atmospheric O_2_. Chemotherapy drugs and inhibitors were purchased in powder form and reconstituted following manufacturers recommendations. Cisplatin (CDDP, Selleckchem), mafosfamide (MAFO; CTX-analog, US Biological), doxorubicin (DOX, Selleckchem), and etoposide (ETO, Merck Millipore), and inhibitors AZD7762 (Selleckchem), CCT241533 (TOCRIS), LY2606368 (Selleckchem), and PF477736 (TOCRIS). Stock solutions for all compounds were prepared in DMSO except for CDDP which was dissolved in DMF (as recommended by supplier). DMSO and DMF were used as vehicle controls and their resulting concentration in culture media was less than 0.1%. We used MAFO in *ex vivo* culture to imitate the *in vivo* activity of CTX, which needs to be metabolized by the liver to generate metabolites with alkylating properties (37,42). During drug treatments, ovaries were cultured with drugs for 2 days, then cultured in drug-free medium for 5 days before further experimentation (7 days total).

The culture medium was changed every 2 days. During inhibitor treatments, ovaries were cultured with inhibitors for 2 hrs prior to irradiation or drug treatment. Following radiation (0.5Gy), medium was changed to inhibitor-free ~24 hrs after radiation. When ovaries were cultured with drugs, replacement to normal medium (drug and inhibitor-free) occurred at 48 hrs after treatment start. Ovaries were cultured for a total of 7 days for oocyte survival analyses or for 3-24 hrs for protein analyses. Drug and inhibitor concentrations used were as follows: CDDP 0.1μg/ml (0.3 μM), 0.25 μg/ml (0.8 μM), 0.5 μg/ml (1.6 μM); MAFO 0.1 μg/ml (0.4 μM), 0.5 μg/ml (1.2 μM), 1 μg/ml (2.4 μM); DOX 25 ng/ml (43.1 nM), 50 ng/ml (86.2 nM), 100 ng/ml (172.5 nM) ; ETO 50 ng/ml (84.5 nM), 100 ng/ml (170 nM), 500 ng/ml (849.5 nM); AZD7762 (0.1, 1 and 10 μM), CCT241533 (0.05, 0.5 and 5 μM), LY2606368 (0.1, 1 and 10 μM), and PF477736 (0.5, 5 and 50 μM).

#### Immunofluorescence and TUNEL staining

Slides with 5 μm ovarian sections were processed using standard methods. Briefly, sections were deparaffinized and re-dehydrated before antigen retrieval using sodium citrate buffer (10 mM sodium citrate, 0.05% Tween20, pH 6.0). Sections were permeabilized in 0.2% Triton X-100 in PBS and blocked with 10% goat serum for 1hr, before incubation with primary antibodies at 4°C overnight. Secondary antibodies were applied for 1hr and slides were mounted with VectaShield (Vector Laboratories) after Hoechst 33342 (Biotechne) staining. Primary antibodies used in this study were mouse anti-p63 (4A4, Biocare Medical, CM163A), rabbit anti-DDX4 (Abcam, ab13840), mouse anti-γH2AX (Millipore, 05-636), rabbit anti-RAD51 (Abcam, ab176458), rabbit anti-53BP1(Novus NB100-304). Secondary antibodies used were Alexa Fluor (Invitrogen). TUNEL staining for apoptotic cells was performed using DeadEnd Fluorometric system (Promega G3250) according to manufacturer’s recommendations.

#### Whole mount staining of ovaries

Whole mount immunostaining of cultured ovaries was performed as described in (159). Briefly, explanted ovaries attached to the polycarbonate membrane were fixed in 2% PFA at 4°C overnight. Fixed ovaries were washed with 70% ethanol overnight and placed in PBS for at least 4 hrs prior to immunostaining. Ovaries were permeabilized in solution containing 0.2% polyvinyl alcohol (PVA), 0.1%NaBH4, and 1.5% TritonX-100 in PBS for 4 hrs, and incubated in blocking solution (10% goat serum, 3% BSA, 0.15% glycine pH 7.4, 0.1% Triton X-100, 0.2% sodium azide, 100 units/mL penicillin, and 100 ng/mL streptomycin, in PBS) overnight. Primary antibody incubation was performed at room temperature with gentle rocking for 2-4 days. Primary antibodies used were mouse anti-p63 (4A4, Biocare Medical, CM163A) and rabbit anti-DDX4 (Abcam, ab13840). Ovaries were washed with wash solution (0.2% PVA and 0.15% Triton-X 100, in PBS) for 1-2 days with buffer changes (2x daily). Ovaries were next incubated with Alexa Fluor secondary antibodies for 2-3 days followed by washing for 1-2 days with buffer changes.

#### Optical clearing, imaging, and quantification of oocytes

Optical clearing of immunostained ovaries was performed as described in (159). Briefly, ovaries were cleared using ScaleS4(0) (40% D-(-)-sorbitol (w/v), 4 M urea, 10% glycerol, and 20% DMSO, in PBS). ScaleS4(0) solution was refreshed twice daily until tissues became cleared (2-3 days). The membranes with ovaries were mounted using CoverWell Incubation Chamber (Research Products International) and imaged using a Leica DM550 microscope. Images were collected as Z-stacks (5 μm) and used to generate maximum projection image in LAS X software. Oocytes were counted on maximum projection whole ovary images using oocyte markers DDX4 (cytoplasmic staining) and p63 (nuclear staining). Small oocytes with diameter less than 30 μm (delineated by DDX4 staining) were categorized as primordial oocytes and larger than 30 μm were categorized as growing oocytes (corresponding to growing follicles including primary and secondary follicles). Percentage survival was calculated by dividing the mean of the treatment group counts by the mean of the control group counts and multiplying by a factor of 100.

#### Immunoblot

Protein extracts were prepared with RIPA buffer supplemented with protease and phosphatase inhibitors (Sigma-Aldrich) using a minipestle or Bioruptor Pico (Diagenode). Proteins were resolved in 10% acrylamide gel and transferred to nitrocellulose membranes. Primary antibodies used in this study were rabbit anti-p63 (Cell Signaling Technology, 13109), rabbit anti-p53 (Leica, CM5P), mouse anti-γH2AX, rabbit anti-DDX4, and mouse anti-ACTB (GeneTex, GT5512). After incubation with HRP secondary antibody, signals were detected using Luminata Forte/Crescendo Western HRP substrate (Millipore). For probing with multiple antibodies, membranes were stripped by using western blot stripping buffer Restore (Thermo Scientific).

#### Statistics

Statistical analysis was performed using PRISM 9.0 (GraphPad Software). To analyze the difference between more than two independent groups (e.g. vehicle vs drug doses) statistical analysis was performed by one-way ANOVA and the significance was determined by Bonferroni multiple comparison test or Kruskal–Wallis for nonparametric data with Dunn’s multiple comparison test. For pairwise comparisons (e.g. wildtype vs. mutant) significance was determined by *t* test or Mann-Whitney nonparametric test. Values of P < 0.05 were considered statistically significant. Data are presented as means ± SEM.*P < 0.05; **P < 0.01; ***P < 0.001; P < 0.0001; n.s., non-significant.

---

**Figure S1.**
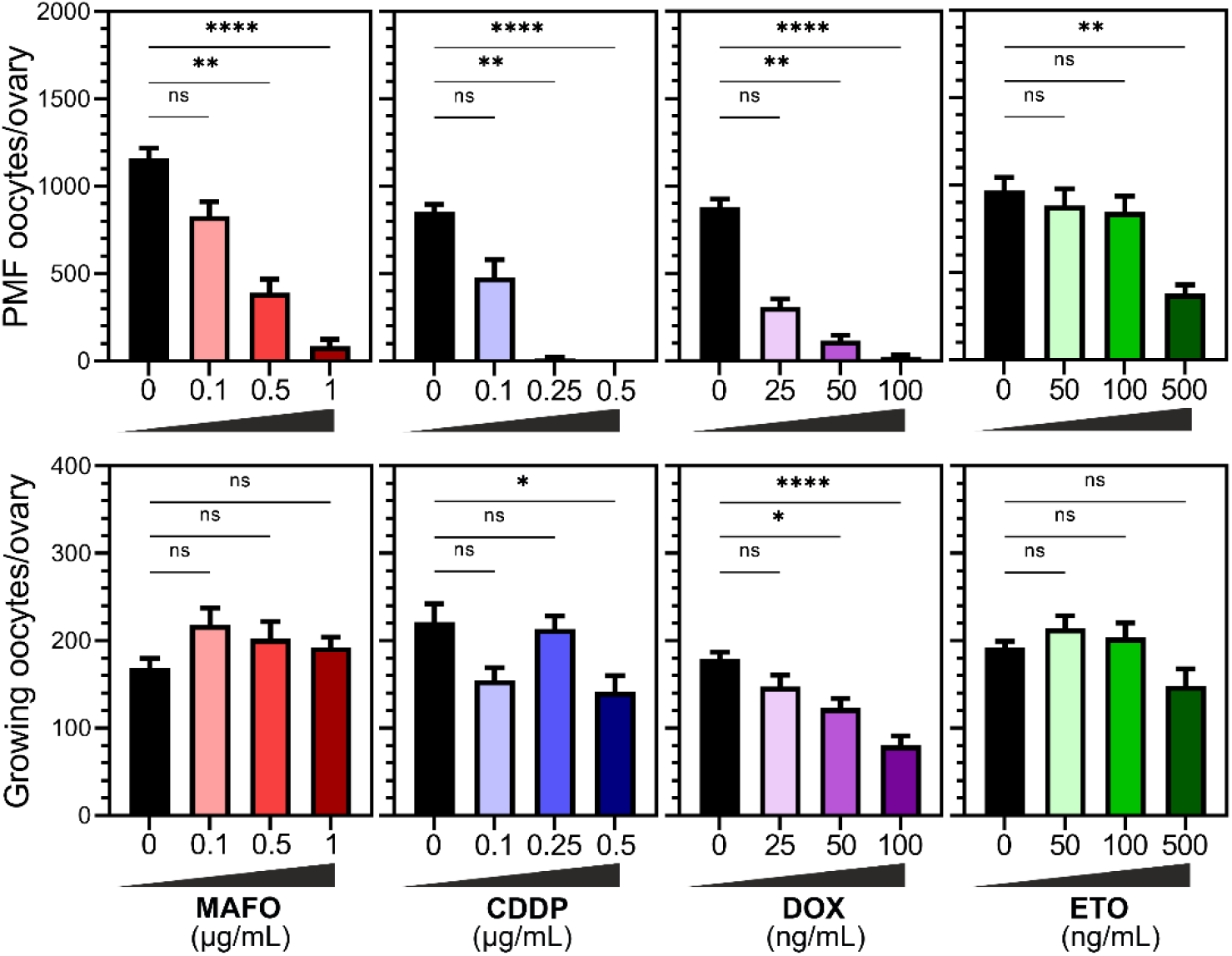
Ovotoxicity of alkylating agents (MAFO and CDDP) and topoisomerase II poisons (DOX and ETO) is dose dependent. Ovaries from wildtype Tg(Pou5f1-EGFP)2Mnn/J females were treated for 48 hrs with MAFO (0.1, 0.5, 1 μg/mL), CDDP (0.1, 0.25, 0.5 μg/mL), DOX (25, 50, 100 ng/mL) or ETO (50, 100, 500 ng/mL). Primordial and growing oocytes were counted in ovaries harvested after 7-day organ culture. Primordial oocytes are found in PMFs while larger growing oocytes are found in primary and secondary follicles present in 2-weeks-old ovaries. Data are expressed as mean ± SEM; *p<0.05, **p<0.01, ***p<0.001, ****p<0.0001 (one-way ANOVA, Kruskal–Wallis with Dunn’s multiple comparison test for nonparametric data).

**Figure S2.**
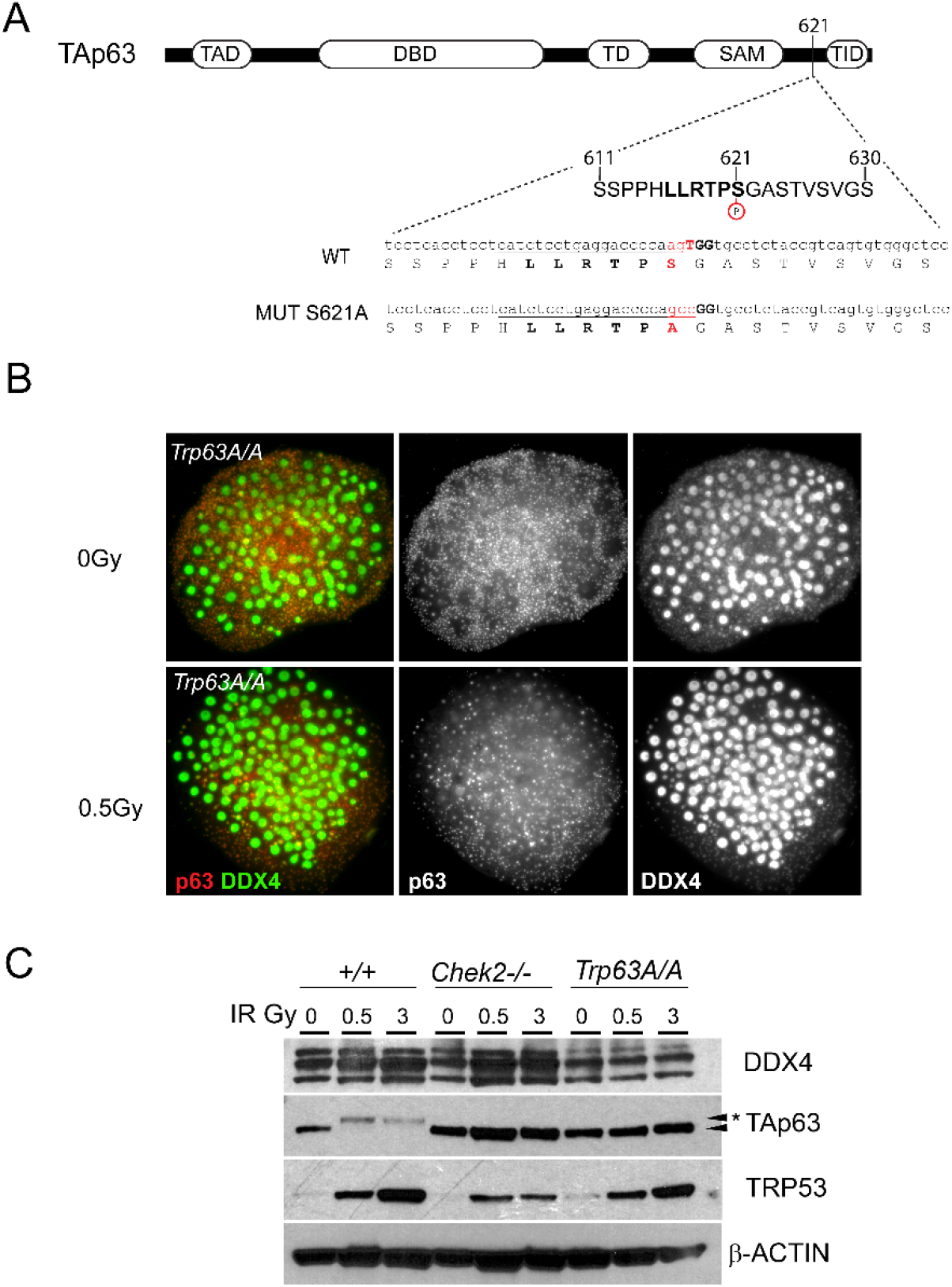
Mutation of 621 Serine to Alanine abolishes TAp63 phosphorylation and leads to its inactivation. (A) Schematic view TAp63 protein domain structure and localization of 621 Serine mutated by CRISPR-Cas9 editing. PAM site is shown in bold and sgRNA sequence is underlined. (B) *Trp63A/A* primordial oocytes show increased resistance to low dose of radiation (0.5Gy). (C) Ovarian protein extracts were collected 3 hrs after radiation with 0.5 and 3Gy from wildtype *Chek2*-/- and *Trp63A/A* ovaries and were analyzed by western blot. TAp63 mobility shift is observed in wildtype ovaries (asterisk) but not in CHEK2 deficient or *Trp63A/A* mutant. TRP53 expression is detected following radiation even in the absence if CHEK2 although at lower level.

